# EEG-to-fMRI Prediction for Neurofeedback: Evaluating Regularized Regression and Clustering Approaches

**DOI:** 10.1101/2025.05.20.654907

**Authors:** Riccardo De Feo, Ekaterina Antipushina, Ivan Tyukin, Yury Koush

**Author notes:** Corresponding authors: R. De Feo,;, Y. Koush,.

## Abstract

While functional Magnetic Resonance Imaging (fMRI) can provide detailed information regarding the functional activity of the whole brain, its cumbersome experimental setting and high cost prevent its application in ecological conditions. This represents a challenge in the context of neurofeedback therapy. To address this challenge one possible approach is to train a predictor for the localized functional activity using multimodal EEG-fMRI records and utilize that predictor in real-time EEG sessions.

Using publicly available multimodal real-time EEG-fMRI data, we present a detailed evaluation of regularized linear regressors both in the context of predicting localized fMRI activity, and to characterize the properties of individual EEG-fMRI runs.

Our results indicate that while it is possible to find clusters of similar time-series in which localized brain activity is highly predictable (Pearson’s r=0.43), the capacity to train regularized linear regressors able to generalize to new subjects remains limited (r=0.24), highlighting two distinct strategies in EEG-fMRI neurofeedback settings.

We characterize the fundamental role for the clustering distance used to identify similar time-series, based both on theoretically-grounded considerations and experimental observations. Our results suggest a clear preference for the cosine metric, instead of the Pearson-based metric utilized in current literature. We further suggest that highly-predictable clusters regular offline paradigms would correspond to high-performing subjects in the context of real-time neurofeedback, and evaluate a regressor-free clustering strategy based on the Log-Euclidean distance of covariance matrices.

## I. Introduction

Functional Magnetic Resonance Imaging (fMRI) can provide a high-resolution, spatially accurate representation of the functional activity of the whole brain. However, besides its limited time resolution, it is also limited by its cumbersome and expensive experimental setup, hindering its application in ecological conditions or in resource-constrained settings. Conversely, while Electroencephalography (EEG) provides a much lower spatial resolution, it is far easier to deploy in a wide range of conditions, at a lower cost, and higher temporal resolution.

The constraints of fMRI are especially relevant in the case of Neurofeedback (NF) [1], where real-time processing is essential and high costs could represent a challenge when scaling NF solutions to clinical application. Furthermore, said constraints represent a challenge in all settings in which ecological or near-ecological conditions would be desirable.

Addressing these challenges, while attempting to retain the informative and spatially accurate fMRI signal where this is not available, requires the introduction of a predictor capable of inferring the fMRI NF signal based on newly-acquired EEG data and EEG-fMRI priors. For this purpose, three methods have been proposed based on classical machine learning. Meir-Hasson et al. [2], [3] proposed to utilize a set of regularized regressors trained on individual EEG runs, and group them together based on the Pearson’s correlation of the model parameters. This corresponds to a cluster of time-series that would then be used to train a common model, referring to this approach as the “EEG Finger-Print” method (EFP). Simoes et al. [4] proposed a method based on random forest regression, while Cury et al. [5] proposed framing the problem as a sparse regression model using spectral features convolved with hemodynamic response functions with different time delays, finding regression coefficient through the FISTA algorithm [6]. Finally, while predictors for fMRI signal from EEG based on deep learning have been developed [7]–[10], it appears that currently none have been successfully applied in NF.

The EFP method has been successfully applied in a number of NF experiments [11]–[17]. Furthermore, regularized regressors are parsimonious with regards to their requirements in training data, and easily interpretable. Because of these characteristics, we believe this method to be of significant scientific interest in the context of NF. However, we found there to be room for improvement with respect to the original choice of clustering methods. Furthermore, while the authors focus on training a regressor on a subset of the available data, the evaluation of this model outside of its cluster and its comparison with a model trained on the whole dataset appeared to be underexplored, as well as the strategy for the selection of the EEG channel utilized to extract the predictive features. As we will show in this work, the use of Pearson’s correlation as a similarity metric between the coefficients of regularized regressors could overlook important information characterizing the EEG data, as the correlations between regressor parameters do not necessarily translate to correlations for their predictions. Based on theoretical considerations and on empirical results, we propose two alternative metrics displaying more desirable properties for the task of clustering regularized regressors, namely the cosine and the Canberra dissimilarities, with a preference for the cosine.

We further analyzed the EFP method based on the choice of clustering parameters, namely the linkage function, the method of comparison between pairs of models, and method of selection of the EEG channels. Our findings indicate that the choice of single linkage and cosine distance (instead of average linkage and Pearson’s correlation) can considerably improve the performance of the regressors within single clusters of EEG runs, reflecting specific modes of behavior within the experimental paradigm. Furthermore, we found an evident trade-off between optimizing for specific clusters or for predictions on any new sample. Finally, our results indicate that a small improvement in the cross-validated prediction outcomes can also be obtained based on considerations concerning the dimensionality of the input dataset.

## II. Methods

### A. Dataset

In the interest of reproducibility we performed all analyses on an openly available dataset, described in detail by Lioi et al. [18], and freely available in the OpenNeuro database [19]. For the sake of completeness we provide here a brief description of the dataset.

#### 1) Data acquisition and pre-processing

The dataset of simultaneous EEG-fMRI data includes 17 subjects. The EEG data were acquired at a 5kHz sampling rate with FCz as the reference electrode and AFz as the ground, using a 64-channel MR compatible Brain Products unit (Brain Products GmbH, Gilching, Germany). The preprocessing included automated gradient artifact correction, downsampling to 200 Hz, and correction for ballistocardiogram artifacts. Concerning fMRI data, they were acquired using echoplanar imaging (EPI) and covering the upper half of the brain on a 3T Siemens Verio MR scanner, with TR=1 s, TE=23 ms, of 2x2x4 mm^3^ voxels and 16 slices, without a slice gap. The regions of interest (ROI) for the calculation of the localized average fMRI NF time-series were selected with an initial functional motor localizer run, identifying the primary motor cortex (M1) and the Supplementary Motor Area (SMA). The NF time-series was computed in relation to the mean signal in a background region unrelated with the NF task (slice 6 of 16), and resampled to a frequency of 4 Hz.

#### 2) Experimental paradigm

The experimental paradigm consisted of 5 EEG-fMRI runs with a 20 s block design alternating rest and task periods (right-hand motor imagery) [19]. Subjects were randomly assigned to two groups based on the provided NF signal (either a sum of EEG and fMRI signals on a single axis or their combination displayed on separate axes). They performed pre-/post-NF motor imagery runs without NF, and three NF runs.

#### 3) Exclusions

For this study, we utilized only the EEG-fMRI NF runs. Two subjects were excluded from our analysis as fMRI NF scores were missing. Furthermore, for one additional subject, the preprocessed EEG signal was not provided for run number 2. Finally, one last subject was excluded because of a mismatch between the length of the NF data and the length of the EEG signal. Due to this, the total number of subjects included was 14, with a total of 41 NF runs.

### B. EFP method reimplementation

#### Features

Following the original formulation of the EFP method [2], [3], we represented the preprocessed EEG signal for each channel as a time-dependent frequency spectrum using the Stockwell transform [20] implemented using the Stockwell python library 1.1.2, with a gamma parameter of 15 and a Kazemi window. The spectrum was initially down-sampled to a frequency resolution of 0.2 Hz and a sampling rate of 4 Hz, and divided into 10 bands up to 60 Hz, corresponding to equal areas on the logarithmic power spectrum. We take as features the average energy within each band. Where the bands were determined for multiple runs at the same time for a common model, they were calculated using the power spectrum of the concatenated signals.

For the prediction of the fMRI NF signal each sample included the last 12 seconds of the power spectrum data, sampled at 4 Hz for each band, resulting in 48 points per band.

#### Models

We used Ridge regressors models [21] as implemented in scikit-learn 1.5.2. When fitting each model, the training data were divided into a learning dataset and a validation dataset using a K-fold split scheme (K = 5) without shuffling the data, to ensure that the validation data originated from intervals temporally distinct from the learning data. The validation data were used to select the regularization parameter alpha from a grid of 50 values, ranging from the minimum of the singular value decomposition of the training data to the square of the maximum singular value. The values were equally spaced on a logarithmic scale, and the chosen alpha depended on the mean squared error metric.

#### Clustering

To train a common model, Meir-Hassen et al. [3] proposed to train a regressor to represent a pattern of behavior that can be identified in a large number of samples. Authors modeled each sample with an individual model (an EEG fingerprint, EFP), and then used a hierarchical clustering algorithm [22] to find the largest cluster sharing the same behavior, based on the coefficients of the regressor models characterizing each sample.

We implemented the clustering algorithm using the Agglomerative Clustering class from scikit-learn 1.5.2, using the average, single and complete linkage methods, and selecting the number of clusters with the elbow method. The choice of average linkage corresponds to the original method.

To calculate the distances between the coefficients of models trained on different band divisions of the EEG spectrum, they were resampled to the same space. This involves both a resampling strategy, and the need to determine a shared set of band boundaries. Originally, resampling was accomplished by first upsampling each set of coefficients to a regular grid, and then averaging them again within the boundaries of the new shared bands [3]. We refer to this interpolation method as MH after the name of the author. As an alternative choice, we contrast this strategy with naive linear interpolation.

To determine the choice of the boundaries for the shared bands, we explored three different strategies. Shared bands were constructed either as shared between all subjects (we refer to this strategy as “Shared”, and corresponds to the original EFP formulation), or between only the two subjects considered when calculating the distances between each pair (“Paired”). We further evaluated the results of this method by directly training every model on the same shared bands from the beginning, including the individual EFPs (“Origin”), eliminating the need for resampling.

Consistently with the original formulation of the EFP method, and to facilitate a fair comparison between the proposed approach and its original implementation, we narrow down the cluster of runs selected for training to the closest 10 from the largest identified cluster.

#### Feature engineering

When experimenting with the dimensionality of the data samples, we utilized the lPCA algorithm implemented in scikit-dimension [23] to estimate the dimensionality of the signal for a given channel, using as input the Stockwell-transformed signal with a frequency resolution of 0.2 Hz. lPCA was fitted using the default parameters.

#### Validation

We evaluated the performance of each according to a leave-one-out (LOO) cross-validation strategy, where one sample was held out from the training process and used for testing, repeating the procedure iteratively for all samples. To evaluate models trained on a single sample, we divided the time series by training on the first 80% of the time series and testing on the remaining 20%. The quality of the regression models was evaluated in terms of Pearson’s correlation with the target NF signal. We focused specifically on correlation as a metric, as in most practical applications the signal could be rescaled and normalized according to the specific experimental paradigms.

### C. Clustering metrics

In this section we discuss the properties of the included dissimilarity measures for the purpose of extracting a relevant cluster of similar data acquisition runs, characterized by a similar activity pattern in the experimental paradigm. We provide a theoretical justification for the use of the cosine similarity [24] or the Canberra distance as clustering metrics [25] in lieu of a Pearson-based dissimilarity. Beyond the dissimilarities discussed below, for the sake of completeness as a general reference, and because of their widespread application, we also included L1 and L2 metrics. However, these are assumed to be overall uninformative for clustering regressors, as they are sensitive to global shifts and scale differences, which may obscure functional similarity.

Let {(*X*^(*i*)^, *y*^(*i*)^), (*X*^(*j*)^, *y*^(*j*)^)} be two independent datasets extracted from two distinct data acquisition runs, where 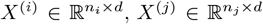, and 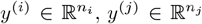.We assume that each dataset is generated by an underlying, unknown and possibly nonlinear function with additive noise:

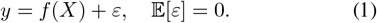

Each dataset is fit independently using Ridge regression with regularization parameters *λ*^(*i*)^, *λ*^(*j*)^ *>* 0:

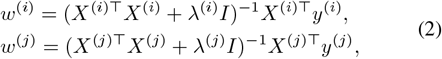

where *I* is the identity matrix.

Let 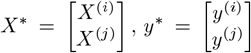 , and let *w*^∗^ denote the Ridge estimator trained on the combined dataset:

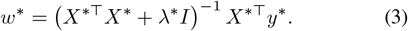

Let us also assume that regularization for the trained regressors sufficiently mitigates overfitting. Each fitted parameter vector *w* can be seen as a regularized linear approximation of *f* . Because we are interested in the correlation with the prediction target, regardless of absolute values, for an ideal dissimilarity metric *d*, for any *X*, we would expect:

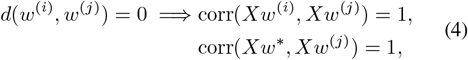

and

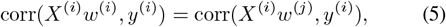

implying that both regressors models similar patterns of behavior in the experimental setting, and that the joint estimator would also model the same behavior. A small dissimilarity between learned weights would also imply functional similarity, and conversely a larger *d* would indicate a degraded performance of the joint estimator, compromising between the misaligned *w*^(*i*)^ and *w*^(*j*)^.

#### 1) Pearson correlation

In the original implementation of the EFP method the measure of distance utilized was based on the Pearson correlation between weight vectors, defined as *d*_*ρ*_(*w*^(*i*)^, *w*^(*j*)^) = 1 − *ρ*(*w*^(*i*)^, *w*^(*j*)^). Explicitly:

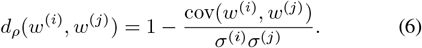

This corresponds to the cosine dissimilarity of the centered vectors:

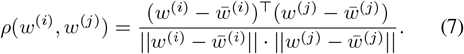

As a consequence, Pearson’s correlation is both shift-and scale-invariant. Its insensitivity to the mean structure of the weights means that *d*_*ρ*_(*w*^(*i*)^, *w*^(*j*)^) = 0 does not imply corr(*Xw*^(*i*)^, *Xw*^(*j*)^) = 1, in contrast with Equation 4.

To see this, note that if *w*^(*j*)^ = *αw*^(*i*)^ + *β***1**, with *α >* 0 and *β* ∈ ℝ, we have *ρ*(*w*^(*i*)^, *w*^(*j*)^) = 1 and *d*_*ρ*_(*w*^(*i*)^, *w*^(*j*)^) = 0. However, in this case the predicted outputs of regressor *w*^(*j*)^ are given by

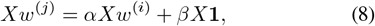

where **1** ∈ ℝ^*d*^ is a vector of ones. The term *X*^(*j*)^**1** sums the feature values across columns for each sample (i.e., it’s the row-wise sum), and thus introduces a structured shift in the predictions of model *j*. In general this does not correspond to a uniform shift, and save for special cases we expect:

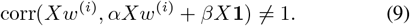

The additive term can introduce a prediction shift capable of degrading the correlation, even when *d*_*ρ*_(*w*^(*i*)^, *w*^(*j*)^) = 0.

#### 2) Cosine dissimilarity

In contrast, the cosine dissimilarity is scale-invariant, but not translation-invariant, defined as:

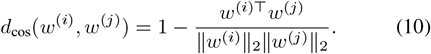

Thus if *d*_cos_(*w*^(*i*)^, *w*^(*j*)^) = 0, we have that *w*^(*j*)^ = *αw*^(*i*)^, resulting in:

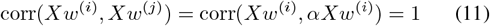

Under the assumption that both datasets reflect similar signal structures and that regularization sufficiently mitigates overfitting, a small cosine dissimilarity between learned weights implies functional similarity between the predictors, consistently with Eq. 4. Cosine dissimilarity between parameter vectors therefore provides a meaningful, scale-invariant proxy for predictive alignment, displaying more consistent behavior than the Pearson correlation between the model weights *w*.

#### 3) Canberra distance

While the cosine distance presents the desirable property of scale invariance, it might overlook fine-grained patterns of considerable importance in the case of NF. In high-dimensional spaces (*d* = 480 in our setting) and especially in the presence of regularization, many components of the weight vector may be small. However those small weights could encode important discriminative behavior characterizing the response patterns of individual subjects. Moreover, in the case of NF, a considerable amount of variance has been observed in the experimental cohorts, suggesting that some features might only be observed in subpopulations [26]. It could then be important to amplify dissimilarities in components where the absolute weight values are small, thus highlighting rare but discriminative features. While in this case the alignment between *w* vectors is not evaluated explicitly, we also experimented with the Canberra distance, defined as:

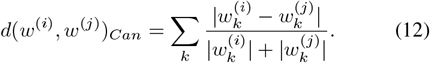

The Canberra distance is not invariant to scaling or translations.

#### Log-Euclidean distance

To provide a metric possibly increasing the practical applicability of the method, by enabling clustering of new samples even in the absence of MRI data, we applied a similarity metric independent of a defined prediction target. For this we utilized the log-Euclidean distance [27] of the covariance matrices of a selection of electrodes. The Log-Euclidean distance has been commonly used to quantify the similarity between positive semi-definite matrices [28] and often used to compare different covariance matrices [29]–[32]. For a pair of symmetric matrices **S**_**1**_, **S**_**2**_ we express this metric as:

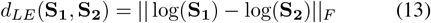

where || · ||_*F*_ refers to the Frobenius norm [33]. To reduce noise and focus on electrodes from a broadly task-relevant region, the selection of channels was limited to the following electrodes: CPz, CP1, CP2, CP3, CP5, P1, P3, P5, Cz, C1, and C3.

### D Channel selection strategies

We compared four channel-selection strategies used to extract the EEG features when training one common model for the entire dataset. Two strategies correspond to selecting the channels where on average we observed the best performance in the individual models, either when trained on individually-defined bands or when using dataset-wide shared bands for that channel.

Alternatively, a similar approach was applied after restricting the selection to the regressors achieving the highest correlation with the NF signal [2]. We set this threshold to include the samples for which at least one regressor for any channel achieved a Pearson’s correlation with the target in the top 25%, calculated globally. Other than observing the application of a similar approach in the existing literature, this is justified by the idea that, assuming non-overfitting regressors attempting to model a possibly highly non-linear function, the joint regressor *w*^∗^ generating predictions 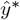 would be expected to achieve lower or equal correlations with the target signal *y*^∗^ than any of the individual models *w*^(*i*)^ with their own individual targets *y*^(*i*)^. In a setting in which the predictability of a specific session or subject could vary significantly, this strategy could better highlight task-relevant channels.

## III. Results

We evaluated the performance of regressors trained on the entire dataset and on subsets of the data, defined by a cluster of similar experimental runs, as a function of the electrode selection strategy, distance and linkage methods, as well as the choice of reference bands and interpolation strategy between the bands. We repeated the same analysis for both available NF targets, referring to the activity in M1 and in SMA.

### A. Channel selection

We evaluated four possible strategies for the selection of the EEG channel utilized to train our models, based on the performance of individual regressors trained on single channels and single EEG-fMRI runs.

#### 1) M1

When the entire dataset was taken into account, the selected electrode was CPz, regardless of band subdivision (Fig. 1). Conversely, limiting the selection to the top 25% EEG-fMRI runs (by best regressor performance on any electrode) pointed to CP1 when utilizing dataset-wide shared bands, and C1 with run-specific bands.

**Fig. 1.**
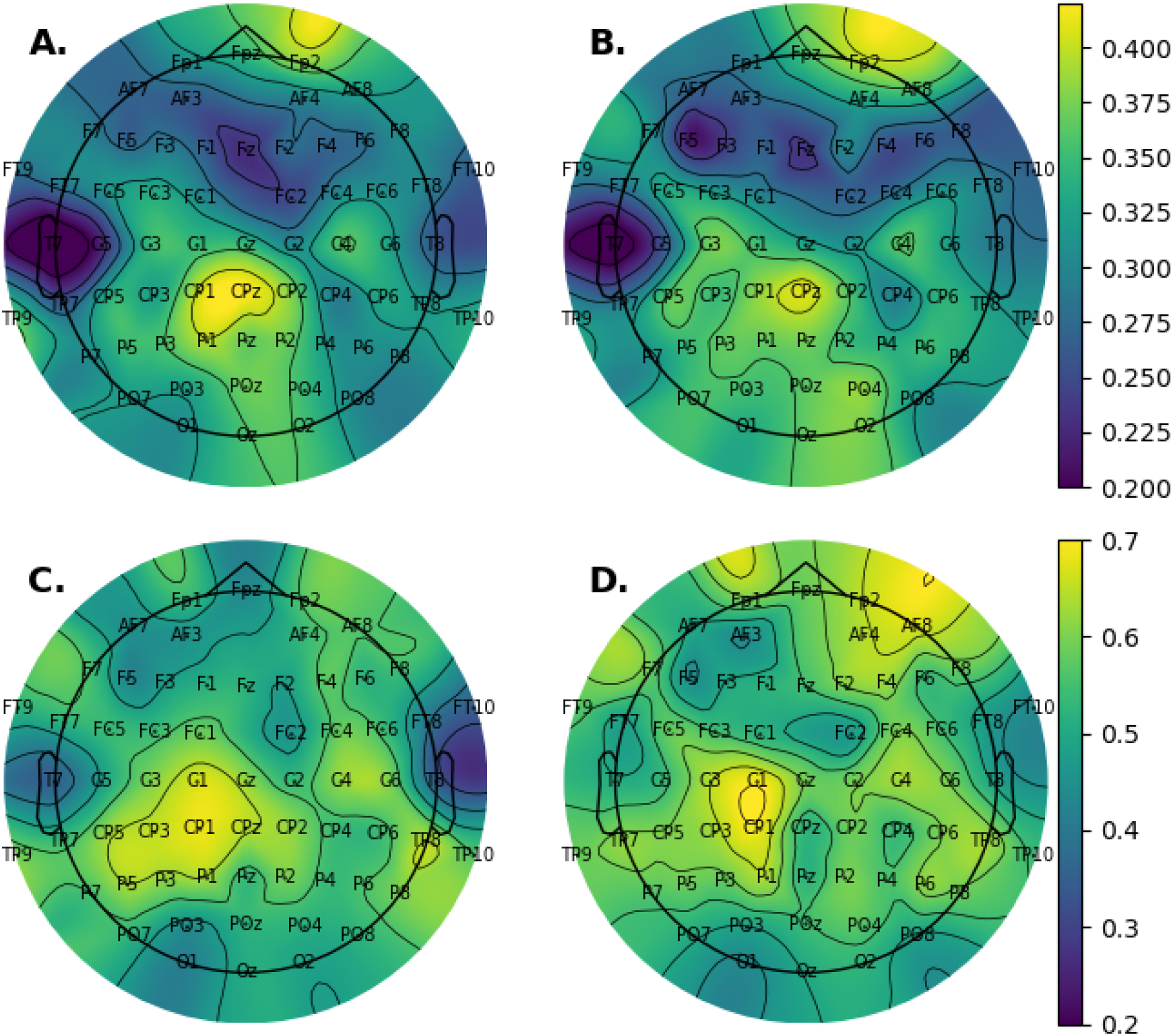
Topomaps of the average Pearson’s correlation by channel for single-channel, single subject models, predicting NF signal in M1. A, B: dataset-wide average; C, D: top 25% runs (any electrode); A, C: models trained on the same dataset-wide bands subdivision; B, D: models trained using run-specific band subdivisions.

#### 2) SMA

All electrode selection strategies point to CPz when using SMA as the regression target, with the exception of selecting only the top 25% runs by correlation while sharing the same bands for all runs, in which case the highest average correlation was observed in CP1 (Fig. 2).

**Fig. 2.**
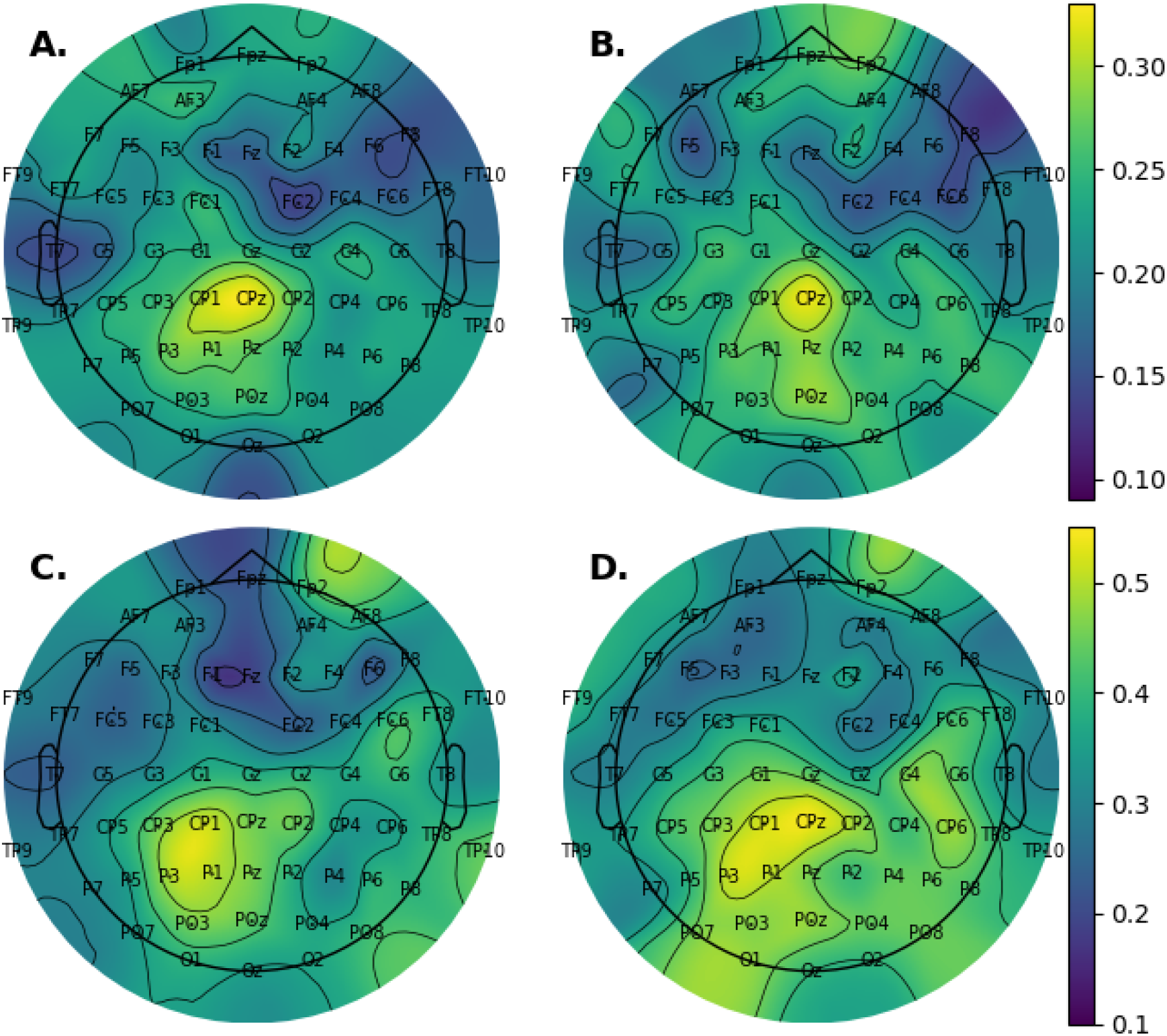
Topomaps of the average Pearson’s correlation by channel for single-channel, single subject models, predicting NF signal in SMA. A, B: dataset-wide average; C, D: top 25% runs (any electrode); A, C: models trained on the same dataset-wide bands subdivision; B, D: models trained using run-specific band subdivisions.

### B. Clustering

We trained a set of models within clusters of experimental runs, and evaluated them inside and outside of their cluster as a function of the clustering parameters, for both of the available prediction targets, M1 and SMA.

#### M1

A detailed breakdown of regressor performance trained on clustered subsets of the data, as a function of all clustering parameters and ROIs, is presented in Figure 3. The best within-cluster performance was observed on the C1 channel using the Canberra distance function and average linkage, comparing the regressors on either Paired or Shared bands, and using MH interpolation, reaching Pearson’s correlation with the target of 0.430. While time-series were clustered individually without giving any weight to the subject’s identity, the cluster of 10 elements thus identified included runs from a total of 5 subjects for Shared bands, and 6 subjects for Paired bands, with a 90% overlap between the two clusters. Although the cosine and Canberra distance functions identified high-performing clusters fairly consistently, this result is not as consistent for Pearson’s distance across different choices of channels, linkage and interpolation strategies.

**Fig. 3.**
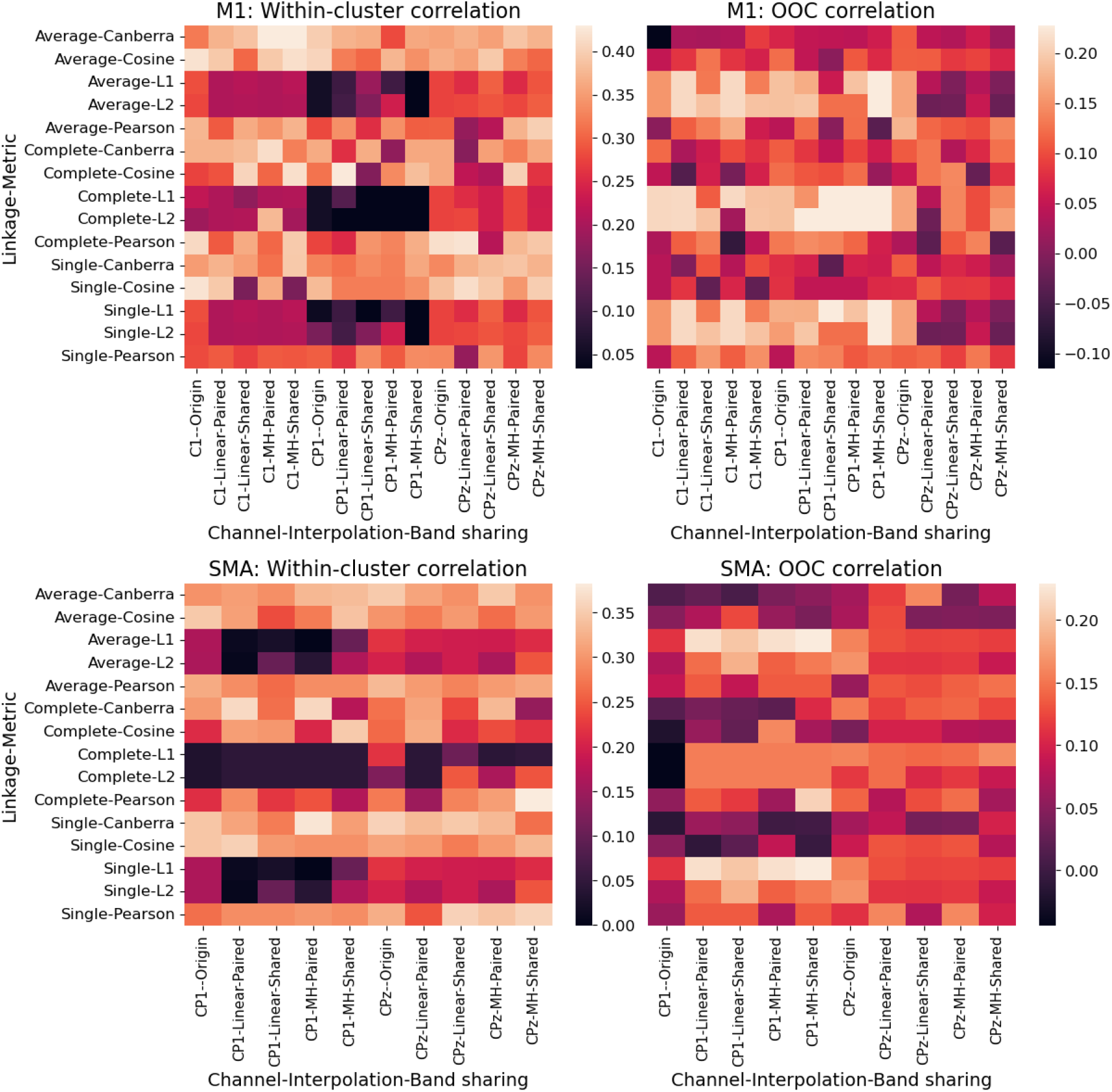
Heatmaps detailing the LOO performance of regressors trained to predict the mean ROI fMRI signal in different clusters, as a function of the clustering parameters, EEG channels, and comparison strategies between regressors. M1: Primary motor area; SMA: supplementary motor area (SMA); OOC: Out-Of-Cluster performance. Linear and MH refer to the interpolation strategy; Origin, Paired, and Shared refer to the choice of bands, as detailed in section II-B.

The original formulation of this method, based on Pearson’s distance and average linkage, achieved correlations with the target signal of 0.399, 0.372 and 0.296 for CPz, C1, and CP1, respectively.

While L1 and L2 distances resulted in a low within-cluster performance, these also corresponded to the highest Out-Of-Cluster (OOC) performance, with a correlation of 0.228 with multiple choices of parameters (Fig. 3). Conversely, the OOC performance of the models previously identified as achieving the best within-cluster performance does not transfer effectively out-of-cluster, with a correlation of 0.088. The Pearson-based, average linkage models described above also appear to be rather uninformative in the OOC setting, with average correlations of 0.141, 0.058, and -0.036, respectively for CPz, C1, and CP1, respectively.

#### SMA

For the SMA target we identify similar patterns concerning the choice of clustering parameters (Fig. 3). The highest results in terms of correlation were consistently achieved using the Canberra and the cosine distances, with the best results achieved using the Canberra distance, single linkage, and paired comparisons (Pearson’s r = 0.378), for the C1 channel. This cluster also naturally grouped together 10 runs from 5 different subjects. As an exception to this pattern, we find a slightly larger within-cluster correlation for the CPz channel using Pearson distance and complete linkage, MH interpolation and shared bands, with a 0.383 correlaton with the prediction target, and including a total of 10 runs from 7 distinct subjects. The original formulation of the EFP method here achieved correlations of 0.317 and 0.292 respectively using the CPz and CP1 channels.

The best OOC results were obtained using the L1 distance, average or single linkage, and the CP1 channel, with a generally low sensitivity to other clustering parameters save for a preference in favor of using any kind of interpolation-based method for comparisons between bands. The highest OOC correlation was 0.229.

#### General patterns of within-cluster performance

While noisy, the general pattern of within-cluster performance suggests two main strategies, consistent across different channels: Cosine distance in combination with single linkage

while strictly using the same frequency bands through the whole process, or the re-implementation of the published EFP method while replacing the Pearson-based dissimilarity with the Canberra distance. Both strategies seem to provide a clear improvement over the original EFP formulation (Fig. 4), either outperforming or matching its performance across all selected electrodes. We also observe that clusters identified by the Log-Euclidean metric appear to be functionally relevant for the M1 target. While these generally underperform compared to regressor-based method, their performance is on par with the original EFP method for channel CP1.

**Fig. 4.**
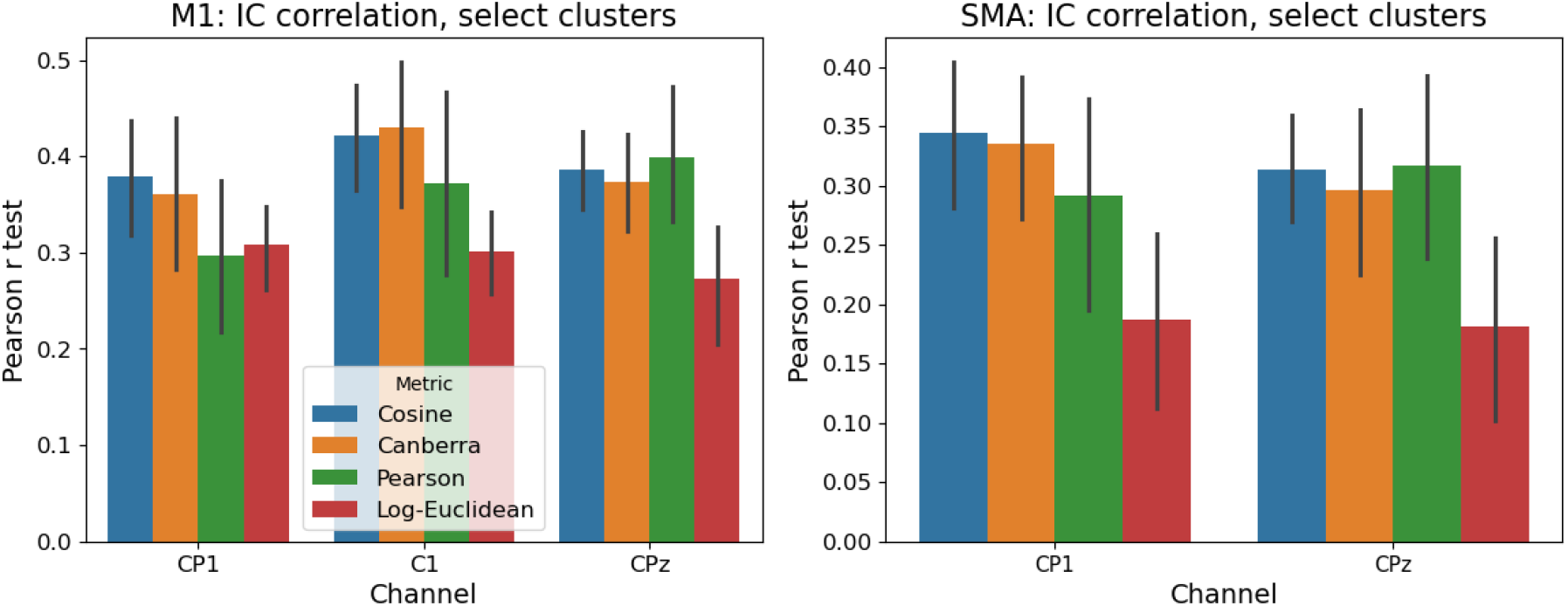
Average performance of models trained and evaluated on within-cluster, using the most reliable clustering parameters for both targets, evaluated via a Leave-One-Out (LOO) validation strategy. The choices of parameters were as follows: Cosine: “Origin” band comparisons, single linkage; Canberra, Pearson: “Shared” band comparisons MH interpolation, average linkage. Pearson corresponds to the original EFP method.

### C. General models

We trained and evaluated regressors using the entire dataset and a LOO strategy, iteratively testing on one subject and using the rest of the data for training. We repeated the procedure for the electrodes designated by each selection strategy. The highest correlation was achieved using channel CPz for both prediction targets, corresponding to the channel with the highest average performance across all individual models, sharing the same bands (Fig. 5). This resulted in a correlation of 0.242 in M1 and 0.202 in SMA.

**Fig. 5.**
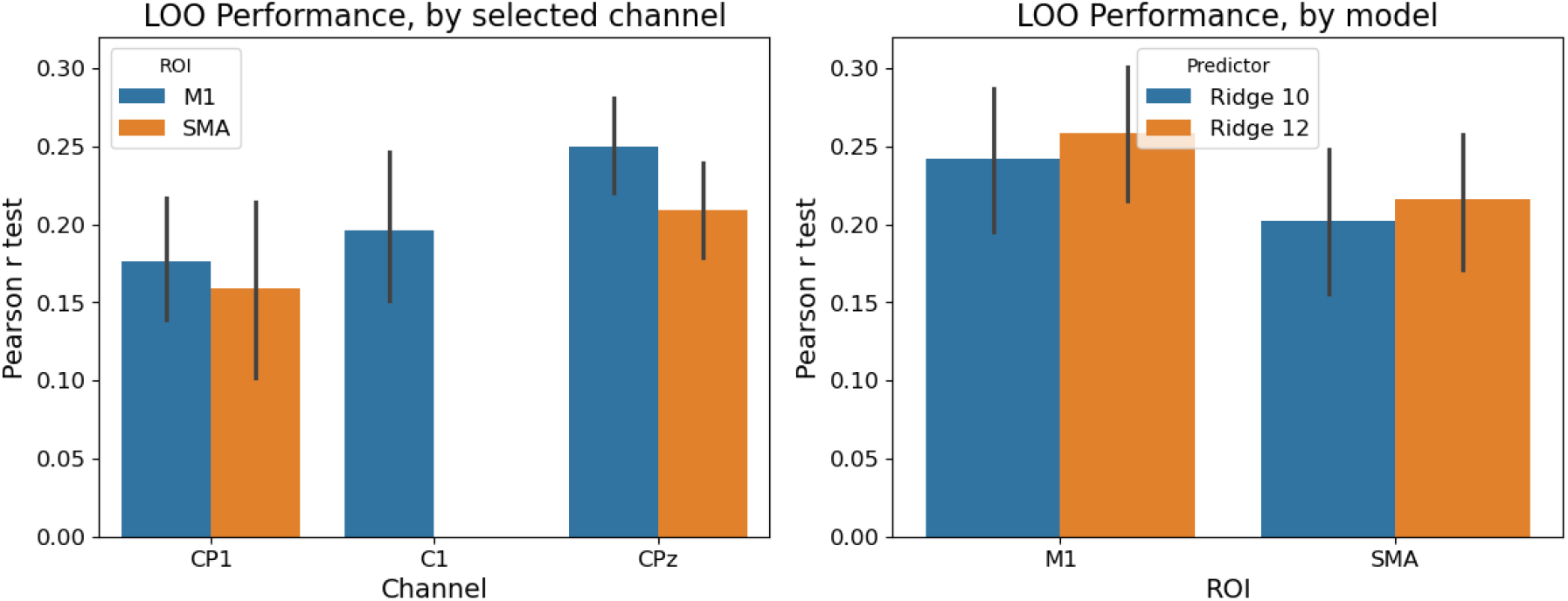
Average performance of models trained on the entire dataset, evaluated via a Leave-One-Out (LOO) validation strategy. Left: LOO correlation of regularized linear regressors (10 bands), by EEG channel of choice and target ROI. Right: LOO correlation of models trained on the CPz channel, by number of frequency bands utilized to characterized each sample.

Using the lPCA algorithm the dimensionality of the signal in the CPz channel was estimated to be 12. We trained a new regressor on new features obtained by subdividing the frequency spectrum in 12 bands rather than 10, resulting in a performance improvement to 0.258 in M1 and 0.216 in SMA, roughly a 7% improvement in both cases (*p <* 0.05, paired permutation test).

### D. Model coefficients and interpretability

The coefficients of the trained regressors characterize the general pattern of localized activity captured by the EEG-fMRI runs used to train them, through their contribution to the predicted NF signal. Figure 6 exemplifies the coefficients for two regressors both trained on the C1 channel, one trained to predict the NF signal from all subjects, and the other exclusively on the high-performing within-cluster runs previously identified.

**Fig. 6.**
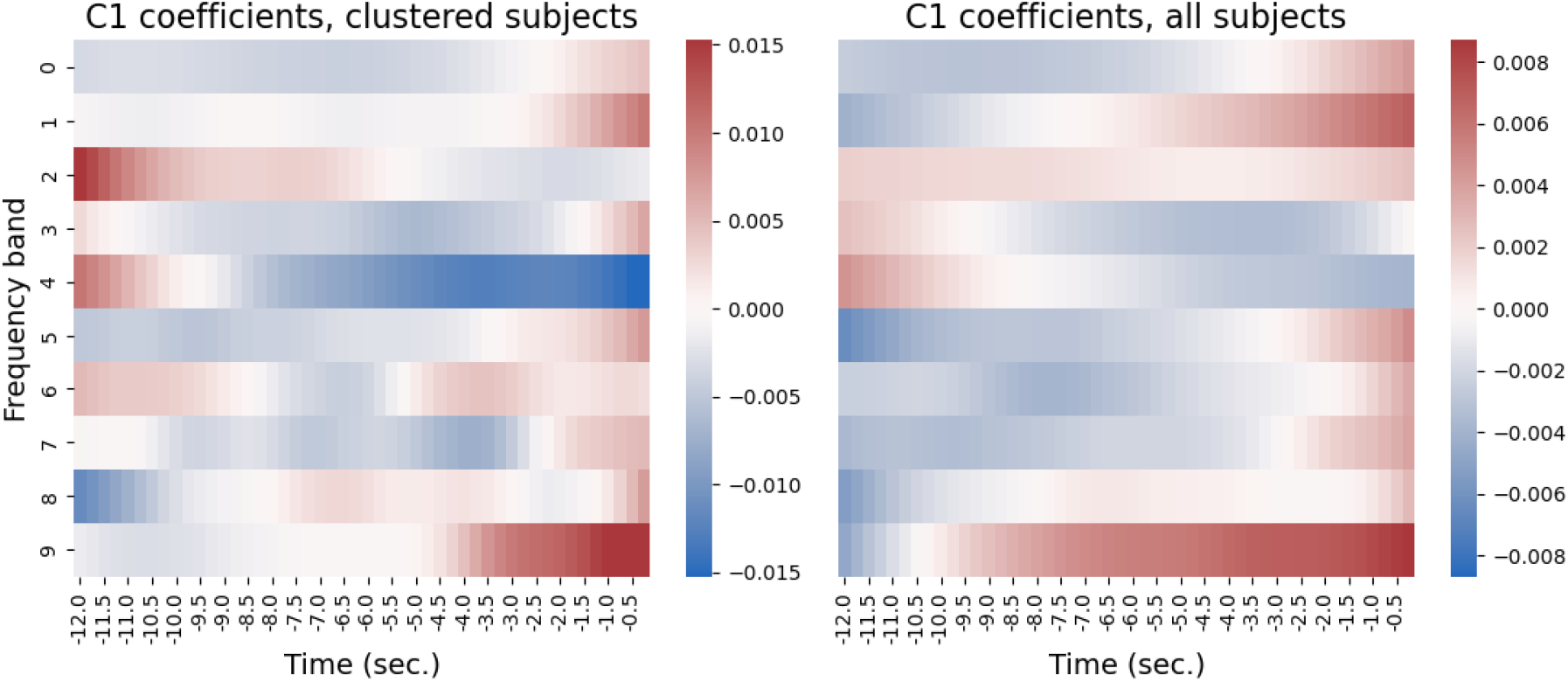
Regressor coefficients for the Ridge regressors trained on channel C1, on a cluster of highly-similar runs (Pearson’s r = 0.430) and across all subjects (Pearson’s r = 0.196). Coefficients are arranged correspondingly to their respective frequency bands (ordinate) and position in time (abscissa).

Qualitatively, the model trained on all subjects displays a smoother distribution of the weights attributed to the signal across time. Conversely, the model trained on the high-correlation clusters displays a more fine-grained time-dependence within each band, subdivided in shorter, smooth and regular blocks, as well as a larger magnitude of the coefficients.

On average, we find that for single-subject, single-channel regressors, the magnitude of the mean of the coefficient vectors *w*, measured in units of their standard deviations 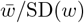, was on average ≈ 0.35.

## IV. Discussion

In our experiments we explored different strategies for training regularized linear regressors to predict the fMRI NF signal in EEG-fMRI experiments. Our findings point to two different strategies to tackle the same problem. One is focused on maximizing the performance on a subset of subjects or sessions, sacrificing the expected performance outside of this subset. We further explored this setting to test the hypothesis that a translation-invariant dissimilarity based on the weights of a regressor would under-perform for this task. The other strategy aims to maximize the expected performance on any subject on average, without focusing on any specific subgroup. In the former case, in line with [3], an effective approach appears to begin by selecting the EEG channels by first only considering those runs in which regressors trained on single channels achieve the highest correlation with the target signal on any electrode, and then selecting the channels displaying the highest average correlation in this subsample. This initial selection is only performed for the purpose of channel selection, and the downstream clustering algorithm is allowed to optimize over the entire dataset. Results concerning the choice of bands on which to train these models were less conclusive. While for the M1 target it was more advantageous to select the electrode by using models trained on individually-selected bands for each run, for the SMA target this was obtained for models trained on the same bands. In most practical settings, however, when training such regressors during the preliminary phases of setting up an EGG-fMRI experiment, comparing

both approaches to electrode selection is inexpensive.

To formulate recommendations as to which methods would be preferable, we looked at both consistency across regression targets and channels, as an indicator of reliability and resilience to both the noise specific to the available dataset, as well as possible variations in terms of tasks and experimental setting. Together with the theoretical considerations of section II-C and its greater simplicity in terms of parameters, avoiding band interpolations altogether, the most reliable measure of dissimilarity appears to be the cosine distance, in combination with single linkage, while training and comparing all models on the same, dataset-wide band subdivision. The drop-in replacement of the Canberra distance in lieu of the Pearson dissimilarity in the original EFP method also yielded comparable results, however the greater degree of complexity of this approach would suggest a preference for the cosine-based method.

By taking into considerations the coefficients of the fitted models, we evidenced that the mean of these vectors was in general non-negligible, and further supports the choice of avoiding a translation-invariant dissimilarity for clustering regressors. As we have shown, a translation invariance in the correlation of the weight vectors of the regressors does not translate to an invariance in the correlations for their predictions.

The individuation of a cluster of highly-predictable, similar runs, mainly from the same subjects, could have different implications from the point of view of neurofeedback. It has been shown that while understudied, NF studies are generally characterized by a heterogeneity in learning and response to the therapeutic approach, with large individual differences in learning performance [26], [34]. It could be possible that subjects characterized by a similar, more predictable response to the paradigm would also be the subjects that better respond to the paradigm itself, perhaps simply as a result of better adherence to the study protocol. In other words, it could be the case that subjects achieving high performance on NF tasks would be reacting in a similar way to the study protocol and thus cluster together, while those who do not might be both more dissimilar, and less predictable. Furthermore, where this regressor-based method has been applied for EEG-fMRI NF (based on Pearson’s correlation as a clustering pseudo-distance), significant improvements have been observed in patient populations [3], [12], [14]–[17]. Considering that the predictive value of this model is reduced for behavior significantly different from the one represented in the training cluster, this suggests that there was a significant overlap between highly-predictable subject and high-performing subjects, and could hint towards the hypothesis that highly-predictable clusters could also in general constitute clusters of high-performing learners. The latter observation could be crucial for cross-modal predictions of fMRI targets based on current real-time EEG signals and EEG-fMRI priors.

These considerations would also suggest the viability of regressor-independent metrics for the identification of high-performing clusters. Limited to our analysis, utilizing the Log-Euclidean distance between covariance matrices we could identify a cluster in which we could train predictors as effective as those trained based on the EFP method, however this was only true for the CP1 channel and M1 target. It is possible that a more thorough exploration in terms of the channels used to build the covariance matrices or by improving the models beyond regularized linear regressors could yield better results in this direction. More generally, this clustering approach could be used to evidence different modalities of response to an experimental paradigm in an unsupervised manner, and to select populations based on the expected response to the paradigm.

As an alternative to the clustering-based approach, it is possible to prioritize the overall performance on any new subject. In this case the straightforward approach of training a regressor on the entire dataset appears to be the best choice, and the choice of channel intuitively points to the selection of the one in which the initial regressors achieve on average the highest correlation, with no initial pre-selection. While in a selection of cases the average OOC correlation was slightly higher for models trained on specific clusters, this improvement is small enough and inconsistent enough that it would be considerably more likely to be a result of randomness rather than a reliable improvement over the naive approach. Consistently, such improvements were not statistically significant. Utilizing lPCA to characterize the dimensionality of the signal and reduce the uncertainty in the selection of the number of bands could instead improve the observed results by a small but statistically significant margin. However we should emphasize that in our approach this was only utilized to select their number, while the bands themselves were still identified only based on the logarithmic power spectrum. Overall such improvements are limited, and with respect to generality it would be more valuable to abandon regularized linear regressors entirely. More promising, recently published approaches based on deep neural network report higher scores for inter-subject evaluations, without focusing on any subset of subjects or sessions [35]. It should be noted, however, that within-cluster LOO correlations for the much simpler linear regressors evaluated in this setting are comparable with the scores currently reported by SOTA DNNs.

This study presents several limitations, owed primarily to limitations in the amount of available data. Because of the paucity of data, we preferred to avoid exploring the possible combinations of multiple electrodes when training and clustering regularized regressors, and could not evaluate different prediction targets. To keep the number of possible parameters configurations under control, we also did not to engage in a systematic exploration of different parameters for the Stockwell transform, or replace it with alternative approaches. However, working with freely-available data ensures that both the data and the code (shared at https://github.com/Hierakonpolis/EEG-fMRI-Regressors) are freely available to all who might be interested.

## V. Conclusions

In contrast with the current approach based on Pearson’s correlation and average linkage, our findings point to the following recommendations when training Ridge regressors on clusters of time-series to predict the mean fMRI signal in a given ROI from EEG data:

- Pre-select channels based on the average performance in the top 25% performers by Pearsons’s correlation, between all runs;
- Use hierarchical clustering with single linkage;
- Evaluate the similarity based on the cosine distance on dataset-wide shared bands.

Alternatively, when trying to train the best general model for any new subject, train on all runs.

Our findings further suggest the hypothesis that clustering-based, data-driven methods could help predict which subjects would better respond to NF therapy, and could be utilized to detect different modalities of response to the NF protocol in an unsupervised manner.

## VI. Contribution statement

RDF: formal analysis, data curation, conceptualization, investigation, methodology, software, visualization, writing. KA: investigation, validation, writing - editing. IT: methodology, writing - review. YK: conceptualization, funding acquisition, methodology, supervision, writing.

